# Enhanced differentiation of functional human T cells in NSGW41 mice with tissue-specific expression of human interleukin-7

**DOI:** 10.1101/2020.04.24.060319

**Authors:** Emilie Coppin, Bala Sai Sundarasetty, Susann Rahmig, Jonas Blume, Nikita A. Verheyden, Franz Bahlmann, Sarina Ravens, Undine Schubert, Janine Schmid, Stefan Ludwig, Constantin von Kaisenberg, Alexander Platz, Ronald Naumann, Barbara Ludwig, Immo Prinz, Claudia Waskow, Andreas Krueger

## Abstract

Humanized mouse models have become increasingly valuable tools to study human hematopoiesis and infectious diseases. However, human T cell differentiation remains inefficient. We generated mice expressing human interleukin (IL-7), a critical growth and survival factor for T cells, under the control of murine IL-7 regulatory elements. After transfer of human cord blood-derived hematopoietic stem and progenitor cells, transgenic mice on the NSGW41 background, termed NSGW41hIL7, showed elevated and prolonged human cellularity in the thymus while maintaining physiological ratios of thymocyte subsets. As a consequence, numbers of functional human T cells in the periphery were increased without evidence for pathological lymphoproliferation or aberrant expansion of effector or memory-like T cells. We conclude that the novel NSGW41hIL7 strain represents an optimized mouse model for humanization to better understand human T cell differentiation *in vivo* and to generate a human immune system with a better approximation of human lymphocyte ratios.

## Introduction

Humanized mouse models have emerged as indispensable tools for improving our understanding of human hematopoiesis and the human immune system. Typically, they are generated by transplanting human hematopoietic stem and progenitor cells (HSPCs) into mice (for review see ^1^). NSGW41 mice carry the hypomorph W41 allele in the *Kit* gene, harbor the NOD-specific variant of the *Sirpa* gene, are T-, B- and NK-cell deficient based on null mutations in *Prkdc* and *Il2rg* genes, and allow for human donor stem cell engraftment in the absence of preconditioning ^2, 3^. NSGW41 mice that are stably engrafted with human hematopoietic stem cell display continuous human hematopoiesis with increased myeloid ^2, 4^, megakaryocytic and erythroid output compared to irradiated NSG recipient mice ^5, 6^.

The introduction of human genes to overcome selective deficiencies in human hematopoiesis has resulted in further improvement of humanized mouse models. Thus, mice expressing human cytokines, including a combination of SCF, GM-CSF, and IL-3 (NSG-SGM3) or M-CSF, IL-3, GM-CSF, and thrombopoietin (MISTRG) show improved myelopoiesis ^7, 8^. Mice carrying a knock-in of human IL-6 show improved B-cell development and function ^9^.

Efficient differentiation of human T cells remains a challenge in humanized mice and we have focused on interleukin-7 (IL-7) to improve that situation. Patients with mutations in the *IL7RA* gene suffer from profound T–B+NK+ severe combined immunodeficiency ^10^. In mice, loss of the *Il7ra* gene results in combined B and T-cell lymphopenia, pointing towards critical cross-species differences ^11, 12^. *In vitro*, murine (m)IL-7 was 100-fold less potent to expand and differentiate human T-cell progenitors when compared to human (h)IL-7 ^13^. Given its role as key factor for lymphocyte survival and proliferation ^10–13^, unrestricted supply of IL-7 might result in unwanted effects including lymphoma generation ^14^. Further, excessive amounts of mIL-7 limits T cell differentiation by interfering with Notch signaling ^15, 16^. Here, we generated hIL-7 (hIL-7) BAC transgenic NSGW41 mice to improve T cell differentiation in humanized mice while simultaneously avoiding unwanted effects caused by excessive and spatially unrestricted availability of hIL-7.

## Results and Discussion

To generate a mouse model with tissue-specific expression of human (h)IL-7, we inserted cDNA encoding *IL7* into a BAC containing regulatory elements of the murine *Il7* locus (Fig. 1a). BAC transgenic mice were generated on the NOD background, crossed into the NSGW41 strain ^2^ and termed NSGW41hIL7. NSGW41hIL7 mice contain three copies of the BAC transgene and expressed hIL-7 mRNA and protein in BM, spleen and thymus (Fig. 1b,c). Engraftment of human HSPCs and T cell differentiation was determined at indicated time points after transplantation of CD34^+^-enriched cord blood cells into unconditioned NSGW41 or NSGW41hIL7 mice (Fig. 1d). Blood from NSGW41 or NSGW41hIL7 mice contained comparable levels of human hematopoietic cells peaking at 16-18 weeks after transplantation (Fig. 1e). However, beginning at week 16 after transplantation frequencies of T cells among human CD45^+^ cells were significantly increased in NSGW41hIL7 mice compared to NSGW41 recipients. Correspondingly, ratios of T and B cells increased progressively over time, with T cells ultimately becoming the predominant lymphocyte population in blood of NSGW41hIL7 mice (Fig. 1e,f). T/B ratios >1 are also observed in human blood. Consistently, in spleens, human CD45^+^ leukocytes were increased in NSGW41hIL7 mice compared to NSGW41 mice, which was mainly attributable to increased numbers of T cells (Fig. S1a,b). The data indicate that expression of hIL-7 under control of endogenous gene regulatory elements fosters T-cell differentiation in NSGW41hIL7 mice.

**Fig. 1:**
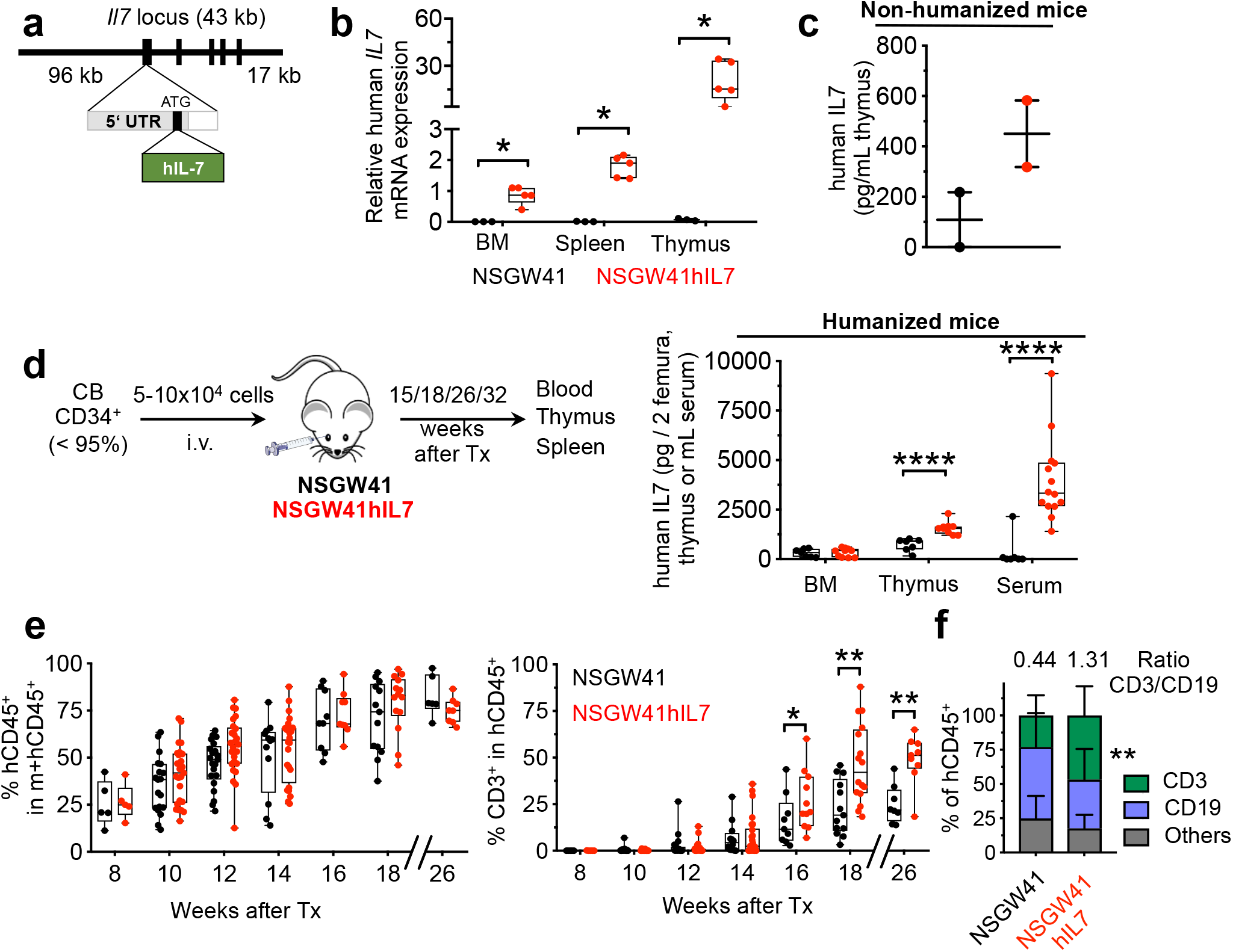
Enhanced generation of human T cells upon transfer of human HSPCs in NSGW41hIL7 mice. **a**, Scheme of BAC construct for the generation of NSGW41hIL7 mice. **b**, Abundance of hIL7 transcript in bone marrow (BM), spleen and thymus from humanized NSGW41 or NSGW41hIL7 mice. Each dot represents an individual mouse. **c**, hIL7 protein levels in thymi isolated from non-humanized NSGW41 or NSGW41hIL7 mice (top) and in bone marrow, thymus and serum from NSGW41 or NSGW41hIL7 mice that have received human HPSCs 26-38 weeks before (bottom). **d**, Scheme of transplantation experiments. **e**, Kinetics of the appearance of human CD45^+^ cells (hCD45^+^, left) and hCD3^+^ T cells within human leukocytes (right) in the blood after humanization. Each dot represents an individual mouse. **f**, Frequencies of T cells, B cells and non-defined other cells of human origin in the blood of NSGW41 (n=13) or NSGW41hIL7 (n=16) mice 18 weeks after humanization. Numbers on top indicate T vs. B cell ratios.

Next, we assessed whether elevated frequencies of peripheral human T cells in NSGW41hIL7 mice were due to more efficient intrathymic T-cell differentiation. Human CD45^+^ cell numbers were 3.3-fold, 3.5-fold, and 21.2-fold higher in thymi from NSGW41hIL7 mice at 15, 18, and 26 weeks after reconstitution, respectively (Fig. 2a). Ratios of human CD4/CD8 double negative (DN), double positive (DP), and CD4 and CD8 single positive (SP) thymocytes were comparable in both recipient lines 15 and 18 weeks after transplantation, indicating that ectopic expression of hIL-7 did not result in aberrant T cell differentiation (Fig. 2b,c). NSGW41hIL7 but not NSGW41 thymi predominantly contained DP thymocytes 26-32 weeks after transplantation. In contrast, DN thymocytes constituted the major population in NSGW41 mice, suggesting that hIL-7 supports human T cell differentiation for extended periods of time in NSGW41hIL7 mice. A recently characterized combined knock-in of hIL-7 and hIL-15 on the NSG background displayed massive skewing towards CD8 SP cells at the expense of DP thymocytes ^17^, suggesting that tissue-specific expression of human cytokines alone is insufficient to promote human T cell differentiation ^18^. Consistent with the T-lineage specific role of hIL-7 in human hematopoiesis, and confirming specificity of transgenic expression of hIL-7, we observed no alterations in B-cell differentiation in NSGW41hIL7 mice compared to NSGW41 (Fig. 2d, S2a,b). We conclude that NSGW41hIL7 mice display improved and extended intrathymic T cell differentiation from human cord blood-derived HSPCs, temporally coinciding with a shift in T/B cell ratios in the periphery.

**Fig. 2:**
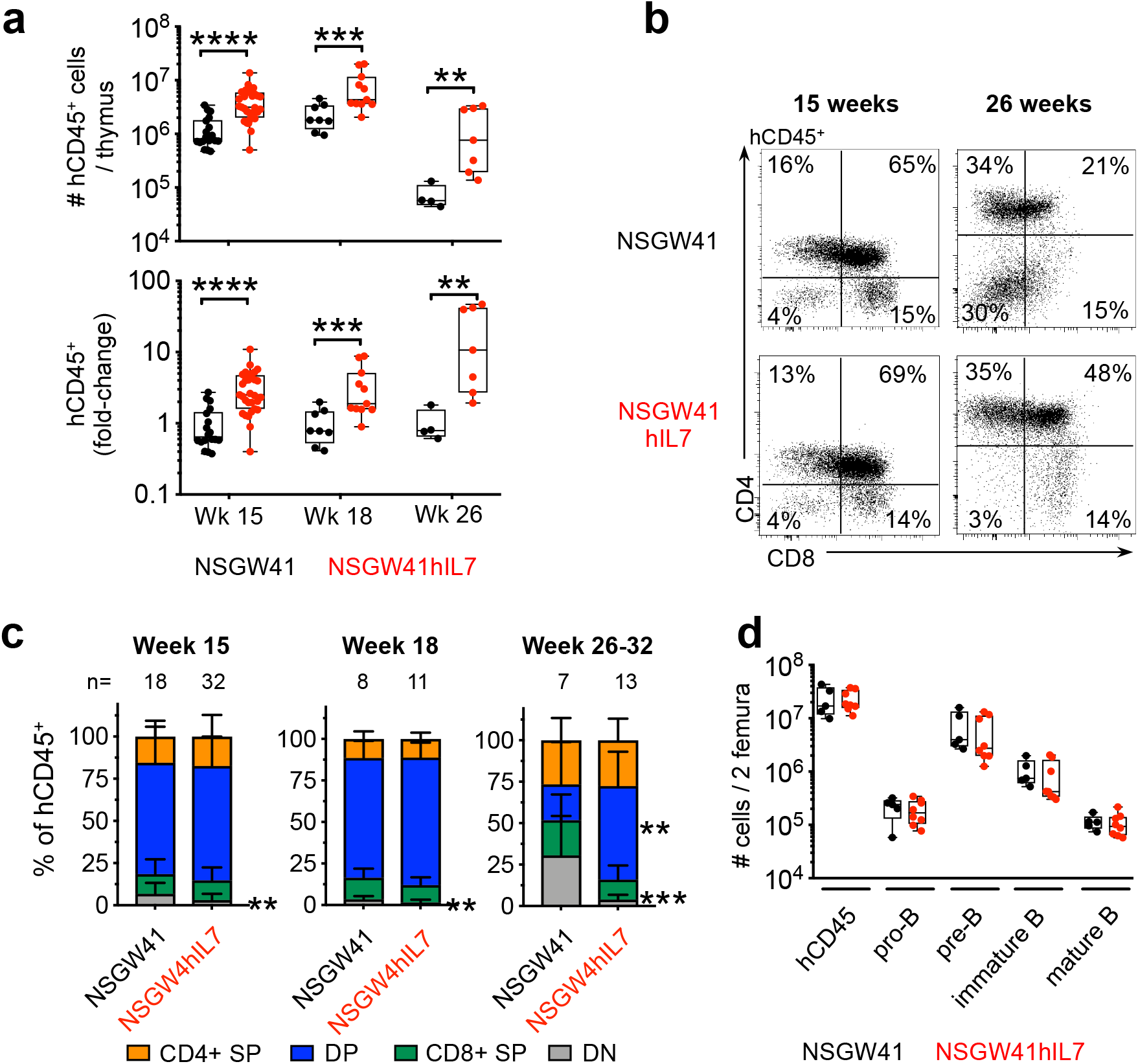
Efficient and prolonged intrathymic T-cell differentiation in humanized NSGW41hIL7 mice. **a**, Numbers (top) and fold-change (bottom) of human CD45^+^ cells in thymi of humanized NSGW41 or NSGW41hIL7 mice at the indicated time points after humanization. Fold-changes were calculated by dividing human CD45^+^ thymocyte numbers from humanized NSGW41hIL7 mice through the thymocyte numbers from humanized NSGW41 mice. This was done separately for each experiment and the results pooled. **b**, Analysis of CD4 and CD8 expression on hCD45^+^ thymocytes from NSGW41 or NSGW41hIL7 mice that have received human HPSCs 15 weeks (left) or 26 (right) weeks before. **c**, Distribution of thymocyte subsets in NSGW41 or NSGW41hIL7 mice at the indicated time points after humanization. **d**, Numbers of B cells subsets in the bone marrow of NSGW41 or NSGW41hIL7 mice 26 weeks after humanization.

Elevated levels of human T-cell precursors in thymus suggested that the increased frequencies of T cells in blood and spleen were largely reflecting increased thymic output. However, hIL-7 might also alter peripheral homeostasis. To test this possibility we further characterized peripheral T cell subsets in NSGW41hIL7 mice. CD4^+^ and CD8^+^ T cell numbers in the blood were increased to comparable extent (Fig. 3a). Within CD4^+^ T cells, frequencies of naive T cells and Recent Thymic Emigrants (RTEs) and effector T cells increased in NSGW41hIL7 mice compared to NSGW41 (Fig. 3b,S3). Concomitantly, frequencies of effector memory T cells were reduced. In CD8^+^ T cells, a similar increase in naive and RTE and decrease in effector memory frequencies was observed, whereas frequencies of T effector and central memory subsets remained comparable to those detected in NSGW41 mice (Fig. 3b, S3). Together, the relative contribution of different subsets to human T cells in NSGW41hIL7 mice resembled the contribution in human peripheral blood. Next, we analyzed T cell receptor (TCR) repertoire diversity using next-generation sequencing to assess possible post-thymic peripheral expansion of T cells. This analysis revealed a high frequency of rare T cell clones, which were comparable between NSGW41 and NSGW41hIL7 mice. Virtually no expanded clones were observed, indicating the absence of hIL-7-induced lymphoproliferation (Fig. 3c). We conclude from these data that elevated peripheral frequencies of human T cells in NSGW41hIL7 mice are largely generated through enhanced intrathymic T-cell differentiation. Furthermore, this route of differentiation results in a composition of the peripheral T-cell compartment largely resembling human peripheral blood. Paucity of peripheral lymph nodes (LN), including mesenteric (m)LN, remains a critical limitation of current NSG-derived humanized mouse models. In NSGW41hIL7 mice mLN were increased in number and individual size compared to NSGW41 mice (Fig. 3d). Given the poor regeneration of lymph nodes in other strains of humanized mice, this observation suggests that NSGW41hIL7 mice might constitute an improved model for studying gut-associated immune responses. Next, we characterized the functionality of conventional human T cells in NSGW41hIL7 mice. To this end, we isolated and activated *ex vivo* CD3^+^ T cells from NSGW41 or NSGW41hIL7 spleens. Upregulation of the bona fide activation markers CD25 and CD69 indicated similar levels of activation (Fig. S4a). Finally, NSGW41hIL7-derived CD4^+^ and CD8^+^ T cells displayed increased divisions in response to anti-CD3/CD28 or phytohemagglutinin stimulation (Fig. 3e, S4). We conclude that human T cells differentiated in NSGW41hIL7 mice respond efficiently to T cell receptor triggering *ex vivo*.

**Fig. 3:**
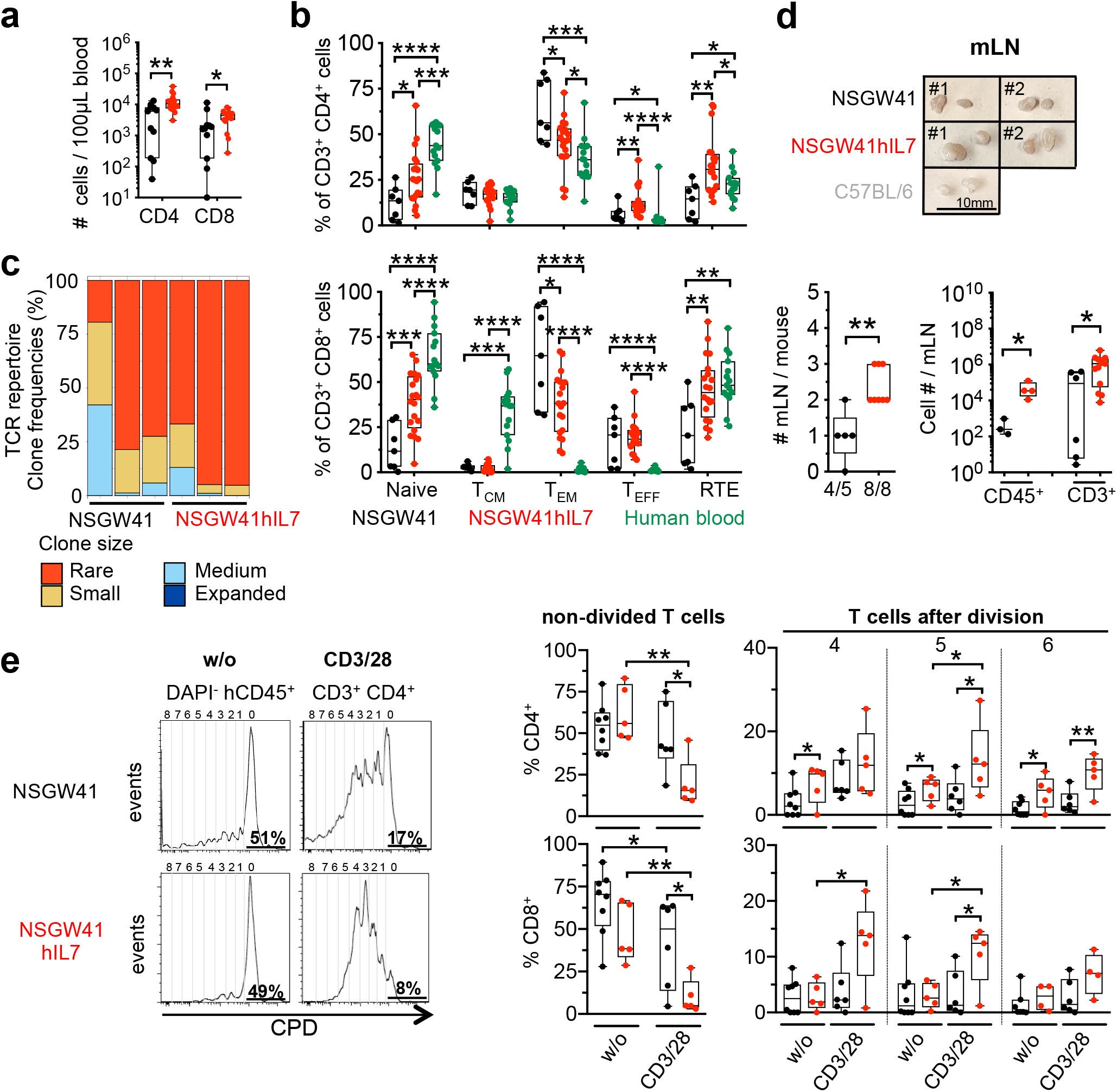
hIL7-BAC transgene increases functional peripheral T cells in the absence of excessive lymphoproliferation. **a**, Human CD4^+^ and CD8^+^ T cells in the blood of humanized NSGW41 or NSGW41hIL7 mice 26 weeks after humanization. **b**, Composition of blood CD4^+^ (top) and CD8^+^ (bottom) T cell subpopulations 26 weeks after humanization: Naïve T cells, central memory (T_CM_), effector memory (T_EM_), T effector (T_EFF_) and recent thymic emigrants (RTE) in NSGW41, NSGW41hIL7 mice, or human blood. **c**, T cell receptor (TCR) repertoire diversity in splenic αβ T cells of NSGW41 or NSGW41hIL7 mice. Clones were binned into rare (0<X≤0.001), small (0.001<X≤0.01), medium (0.01<X≤0.1) and expanded (0.1<X≤1) (n=3 per group). **d**, Photographs of mLN from humanized NSGW41 or NSGW41hIL7 mice isolated 26 weeks after humanization, or C57BL/6 controls (top). mLN number per mouse (bottom left). hCD45^+^ and hCD3^+^ cell numbers in mLN from NSGW41 or NSGW41hIL7 mice isolated 26 weeks after humanization (bottom right). **e**, Activation of human T cells from NSGW41 or NSGW41hIL7 mice. Histograms depict division of CDP-labeled spleen hCD3^+^ T cells 6 days after stimulation with CD3/28 beads or control (w/o, left). Frequencies of non-divided human T cells and T cells that have divided 4, 5 or 6 times 6 days after stimulation with CD3/28 antibody-coated beads or controls (right). Top: CD4^+^ T cells, bottom: CD8^+^ T cells.

Regulatory T (Treg) cells are central in providing protection from autoreactive T cells ^19^. However, in the context of malignancy Treg-cell mediated immunosuppression precludes effective anti-tumor immune responses ^20^. Within the total T cell pool, Treg cells constitute a comparatively small population, making their functional analysis difficult in extant humanized mouse models with low overall T cell numbers. We observed similar frequencies of regulatory T (Treg) cells in multiple organs from NSGW41hIL7 or NSGW41 mice (Fig. 4a). However, given the higher overall T cell numbers in NSGW41hIL7 mice, Treg cell numbers were increased making their analysis feasible (Fig. 4a). To study the activation of Treg cells in humanized NSGW41hIL7 mice, we transplanted porcine pancreatic islets into the portal vein of humanized NSGW41hIL7 mice ^4^. 18 hours after xenotransplantation, Treg cells in the liver but not spleen displayed significantly increased expression of HLA-DR evidencing site-specific activation through the xenograft (Fig. 4b). We conclude that humanized NSGW41hIL7 mice represent a good model for the study of human Treg cells *in vivo*.

**Fig. 4:**
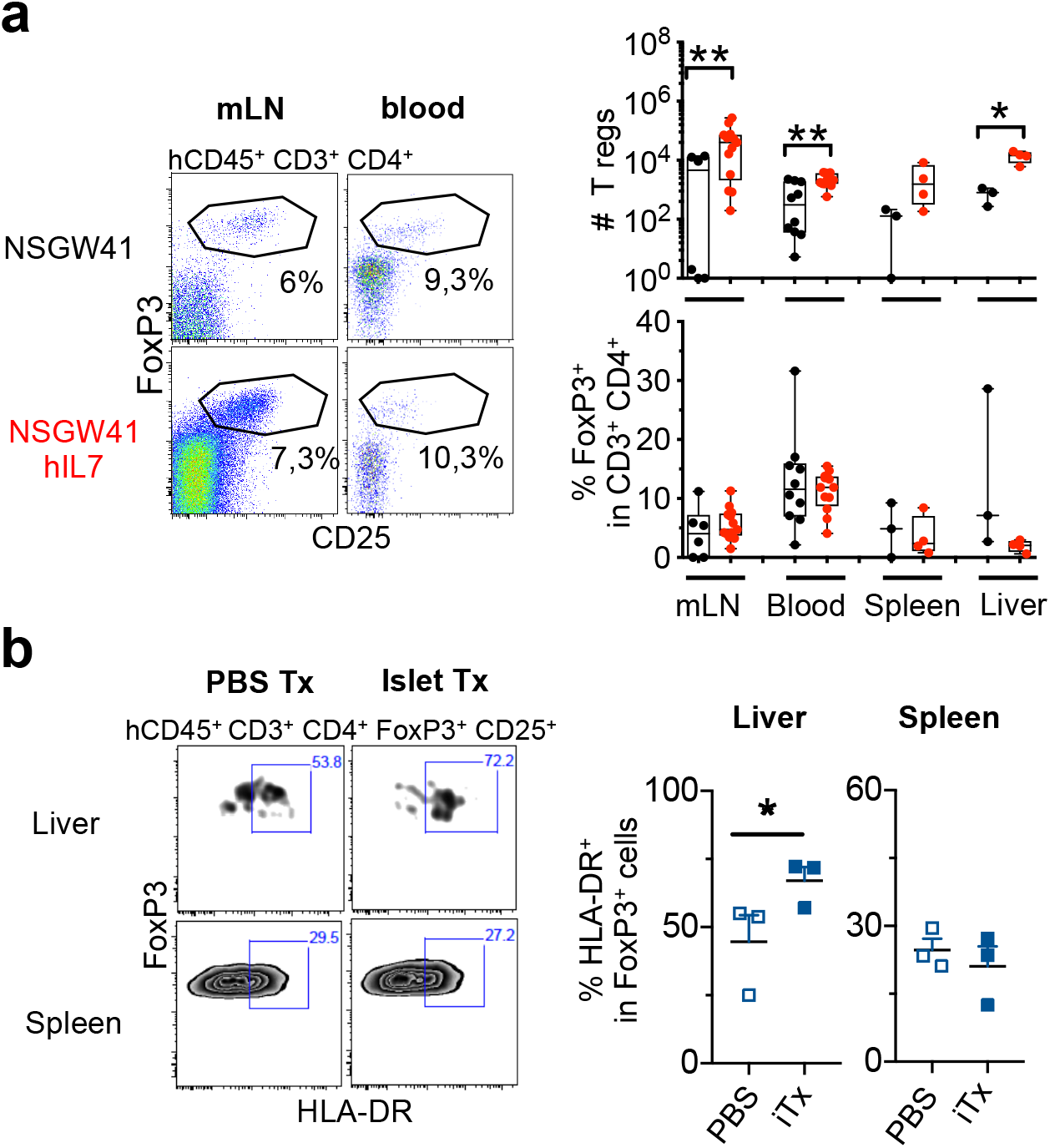
Efficient generation of human Treg cells in NSGW41hIL7 mice. **a**, Representative dot plots analyzing mesenteric lymph nodes (mLN) and blood (left) and numbers (right, top) and frequencies (right, bottom) of Tregs in mLN, blood, spleen and liver of humanized NSGW41 or NSGW41hIL7 mice. **b**, Human activated Tregs after intraportal xenotransplantation of porcine pancreatic islets into NSGW41hIL7 mice 26 weeks after humanization. Representative dot plots of liver and spleen analysis (left). Frequencies of HLA-DR^+^ FoxP3^+^ T cells in spleen and liver of humanized NSGW41hIL7 mice (right) 18 hours after transplantation of islets (iTx) or PBS.

Taken together, we have demonstrated that spatially restricted expression of hIL-7 in NSGW41hIL7 mice results in an improved capacity for T-cell differentiation from human HSPCs. Thus, we observed T/B cell ratios geared towards a higher abundance of T cells when compared to NSGW41 and other humanized mouse models ^21^. Notably, increased intrathymic T-cell development promoted the generation of a peripheral T-cell pool with a high TCR diversity and an overall composition of T-cell subpopulations reminiscent of human peripheral blood. Nevertheless, thymus size and cellularity in humanized NSGW41hIL7 mice remained smaller than that of wild-type mice, suggesting that additional factors, presumably depending on lymphocyte-stromal cell interactions, are required to generate a normal-sized thymic microenvironment after humanization. It has been shown that such a limitation can be partially overcome by co-transplantation of *in vitro*-differentiated proT cells ^22^.

We detected no evidence for hIL-7-driven lymphoproliferation. We designed NSGW41hIL7 mice to express hIL-7 using regulatory elements that allowed to faithfully identify cells expressing mIL-7 in an earlier study ^23^. Our study indicates that this expression pattern and the observed levels of expression of hIL-7 are able to strike a balance to overcome the limited potency of mIL-7 on human lymphocytes, while at the same time avoiding detrimental effects of aberrant expression of hIL-7 ^13, 14^.

Enhanced T-cell differentiation in NSGW41hIL7 mice allowed for analysis of T cell subsets, including Treg cells. In addition, size and number of mLN were increased. Thus, humanized NSGW41hIL7 mice have the potential to foster investigation of rare human T-populations in vivo as well as gut-associated immunity.

## Methods

### Generation of NSGW41hIL7 mice

NODhIL7 mice were generated by pronuclear injection of a BAC containing codon-optimized cDNA of *hIL7* introduced at the 3’ end of the 5’UTR of the *Il7* gene flanked by 96kb upstream, and the entire *Il7* locus plus an additional 17kb downstream. The BAC was constructed according to a described strategy ^23^. BAC integrity upon integration and copy numbers were determined by PCR. Offspring showing detectable expression of hIL-7 mRNA was crossed with NSGW41 mice ^2^ to generate the NSGW41hIL7 strain. All animal experiments were performed in accordance with German animal welfare legislation and were approved by the relevant authorities: Landesdirektion Dresden, the Thüringer Landesamt für Verbraucherschutz, the Niedersächsisches Landesamt für Verbraucherschutz und Lebensmittelsicherheit (LAVES), and the Regierungspräsidium Darmstadt.

### Human HSPC transplantation

Cord blood samples were obtained from the Department Obstetrics, Gynecology and Reproductive Medicine, Hannover Medical School, Hannover, from the Bürgerhospital Frankfurt am Main, and from the DKMS Cord Blood Bank, Dresden, and were used in accordance with the guidelines approved by the ethics committees of Hannover Medical School, Frankfurt University Clinics, Dresden University of Technology, and University Clinics Jena. CD34^+^ HSPCs were isolated using dual magnetic beads enrichment according to the manufacturer’s instructions (Miltenyi Biotech) ^2^. Purities >95% were considered acceptable. Contaminating T cell frequencies were routinely below 1%.

### Flow cytometry

Analysis was performed as described before ^2^. A full list of antibody panels is provided in Table S1.

### TCR repertoire analysis

After mRNA isolation (Qiagen Micro Kit), cDNA was generated via the Smarter 5’RACE cDNA amplification kit (Clontech) using 4.5μl mRNA input and following the recommended protocol. Complementarity-determining region 3 (CDR3) regions of the human TRB locus were amplified through a gene-specific primer (2μM final concentration) that targets the constant region of the beta (β)-chain (GCACACCAGTGTGGCCTTTTGGG) and a primer (1μM final concentration) binding to the introduced SMARTER oligonucleotide (CTAATACGACTCACTATAGGGC) using the Advantage 2 PCR kit (Clontech) in a 50μl reaction. Both primer sequences further contain 16S Illumina overhang adapter sequences. Cycling conditions were as following: 120 s 95°C; 30 times 30 s 95°C, 45 s 64°C, 60 s 72°C; 60 s 72°C. Generated PCR amplicons were agarose gel purified (Qiagen GelExtract.) Next, PCR samples were indexed with Nextera Illumina Indices reads using the Advantage 2 PCR kit (Clontech) in a 8 PCR cycle reaction and purified with Agencourt AMPpure XP beads (Beckman Coulter) according to the manufacturers protocol. Samples were pooled, denatured and subjected to Illumina MiSeq analysis using 500 cycles and paired-end sequencing following Illumina guidelines. Sequencing libraries contained 20% PhIX for library complexity. Demultiplexed Fastq files were annotated to the human TRB locus via MiXCR software ^18^. Individual CDR3 nucleotide sequences were ranked according to their abundance within the respective samples and further analyzed using VDJTools ^24^ and TcR ^25^. TCR repertoire data are available at SRA (https://www.ncbi.nlm.nih.gov/sra), accession number PRJNA606460.

### Islet xenotransplantation

500 IEQ (Islets Equivalent) adult pig islets or PBS were transplanted in the portal vein of NSGW41hIL7 mice that were humanized 24 weeks before. Islets were obtained from Goettingen minipigs (Ellgard) as described before ^26, 27^. Human blood cell chimerism of used mice was 32-85%. Regulatory T cells were analyzed 18 hours after surgery.

### T cell co-stimulation

hCD3^+^ T cells were isolated from the spleen of humanized NSGW41 or NSGW41hIL7 mice 16 to 25 weeks after HSPC transplantation, enriched using negative depletion (Miltenyi), and CPD labeled (eBioscience). 10^5^ hCD3^+^ T cells were mixed with human T-Activator CD3/CD28 Dynabeads (Thermofisher, ratio 3:1) or PHA (1μg/mL) in RPMI 10% FCS, 20mM L-glutamine, 10mM Hepes, 1mM Sodium Pyruvate, 50μM ß-mercaptoethanol with recombinant hIL-2 (30 U/mL) and incubated for 6 days (37°C, 5% CO2).

### Statistics

Student’s t tests were performed for all statistical analyses using Prism 8 for MacOSX software. In all graphs *p = 0.05–0.01, **p = 0.01–0.001, and ***p = 0.001-0.0001 and ****p < 0.0001; data represent the mean ± SD. Boxes and whiskers display the data distribution through their quartiles.

## Acknowledgements

We are grateful to Katrin Witzlau and Esther Imelmann for technical assistance and management of mouse colonies. We would like to thank Dr. Ellen Richie (MD Anderson Cancer Center, TX, USA) for providing BAC constructs and advice on cloning. The work was supported by grants from the German Research Foundation (DFG, SFB738-A7 and SFB902-B15) (to A.K.) and FOR2033-A03, TRR127-A5, WA2837/6-1, WA2837/7-1 and through the project EDISCIDPROG funded by the EU ERA-Net for Research Programmes on Rare Diseases, E-Rare (to C.W.).

## Authorship contributions

E.C. planned, conducted, interpreted experiments and wrote the paper. B.S. planned, conducted and interpreted experiments. S.Rah, J.B., and N.V. conducted experiments. J.B., S.Rav., and I.P. conducted and interpreted experiments on TCR repertoire. U.S., J.S., S.L. and B.L. isolated and provided porcine islets for xenotransplanation. F.B., C.v.K, and A.P. provided crucial reagents. R.N. generated knock-in mice, and C.W. and A.K. conceived the study, planned and interpreted experiments and wrote the paper.

## Disclosure of conflicts of interest

The authors declare that no conflicts of interest exist.

## Supplemental Material

**Table S1.**

| Antibodies | Clone | Provider |
| --- | --- | --- |
| CD3 APC, AF780, APC Cy7 | UCHT1, HIT3a | eBioscience |
| CD4 PE Cy7, Pacific Blue | OKT-4/RPA-T4 | eBioscience |
| CD8 PE Cy5, PerCP, PE Cy7 | SK1, HIT8a | eBioscience |
| CD10 APC Cy7 | HI10a | BioLegend |
| CD14 AF700 | M5E2 | BD Pharmingen |
| CD16 PerCP, PE Cy5 | 3G8 | BioLegend, BD Biosciences |
| CD19 FITC, PC7, APC, FITC | HIB19 | eBioscience |
| CD25 PE | M-A251 | BioLegend, BD Biosciences |
| CD31 PE | MEM-05 | Immunotools |
| CD33 PE, PE Cy7 | WM-53 | eBioscience |
| CD34 PE Cy7, Pacific Blue | OKT-4/RPA-T4 | eBioscience |
| mCD45 AF780, AF700 | RA3-6B2 | eBioscience |
| hCD45 eF450, FITC, PE Cy7, PerCP | HI30 | eBioscience, BioLegend |
| CD45RA eF450 | HI100 | eBioscience |
| CD45RO FITC, PE | UCHL1 | BD Biosciences |
| CD62L Biotin | DREG.55 | eBioscience |
| CD69 PerCP Cy5.5 | Y1/82A | BioLegend |
| CD197 APC, BV510 | REA337,<br>G043H7 | Miltenyi |
| CD235 FITC | GA-R2 (HIR2) | eBioscience |
| Foxp3 APC | PCH101 | eBioscience |
| HLA DR PerCP Cy5.5 | 4S.B3 | BioLegend |
| IgD PE | IA6-2 | BD Pharmingen |
| IgM Biotin | SA-DA4 | BioLegend |
| Ter119 FITC | 11-5921-82 | eBioscience |
| Streptavidin V500, FITC | na | BD Biosciences,<br>BD Pharmingen |
| Cell Proliferation Dye eF450 | na | ebioscience |
| Fixabe viability dye AF700 | na | BD Biosciences |

### Supplementary figures

**Supplementary Figure 1:**
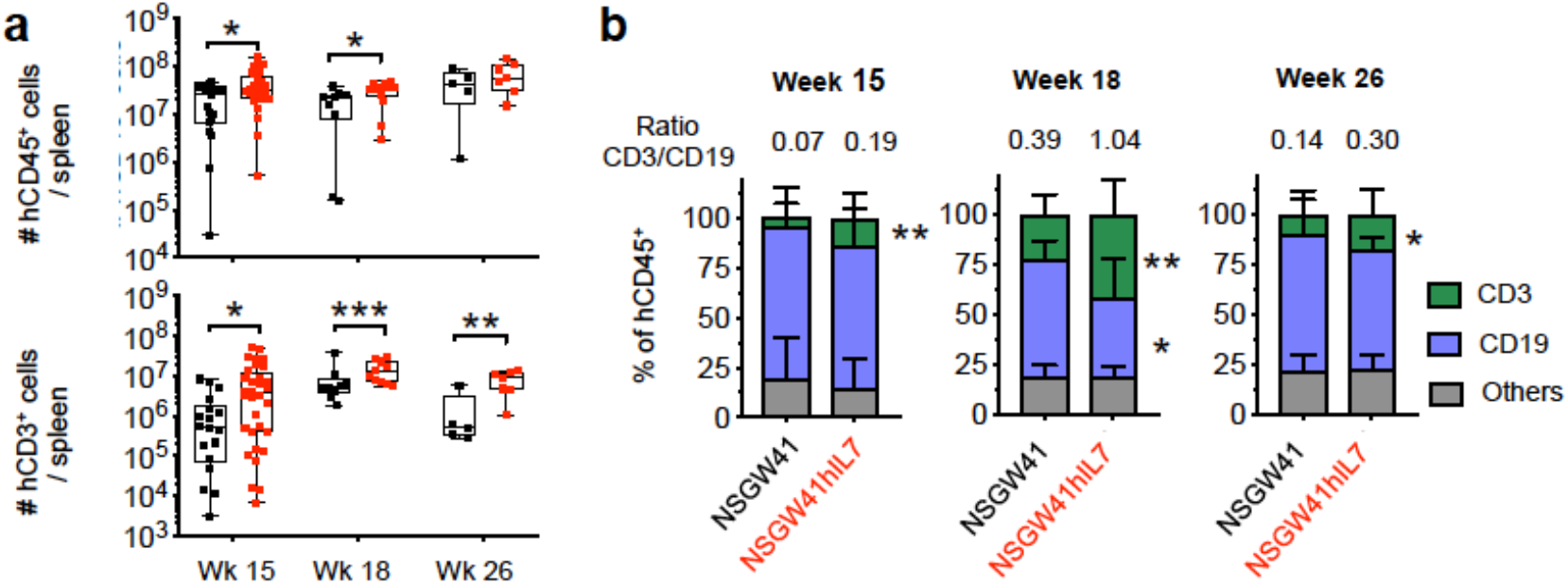
Human lymphocyte composition in spleen. (**a**) Numbers of hCD45^+^ leukocytes (top) and hCD3^+^ T cells (bottom) in spleens of NSGW41 or NSGW41hIL7 mice 15, 18 and 26 weeks after humanization. (**b**) Frequencies of hCD3^+^, hCD19^+^, and other cells within hCD45^+^ leukocytes in spleens of NSGW41 or NSGW41hIL7 mice 15, 18 and 26 weeks after humanization. Numbers on top of graphs indicate T vs. B cell ratio.

**Supplementary Figure 2:**
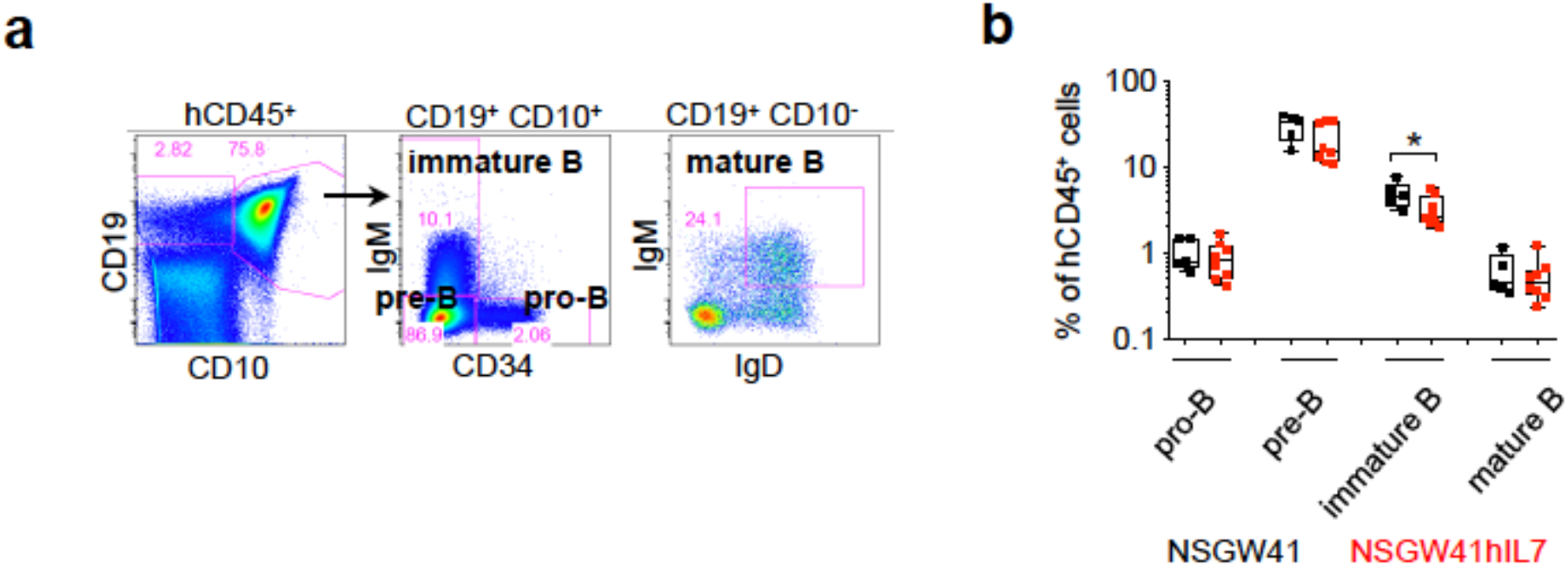
B lymphopoiesis in NSGW41hIL7 mice. (**a**) Dot plots show the gating strategy of bone marrow human B lineage subsets: pro-B (hCD45^+^ CD19^+^ CD10^+^ IgM^−^ CD34^+^), pre-B (hCD45^+^ CD19^+^ CD10^+^ IgM^−^ CD34^−^), immature B (hCD45^+^ CD19^+^ CD10^+^ IgM^+^ CD34^−^) and mature B cells (hCD45^+^ CD19^+^ CD10^−^ IgM^+^ IgD^+^). (**b**) Frequencies of B cell subsets in bone marrow of humanized mice 26 weeks after humanization.

**Supplementary Figure 3:**
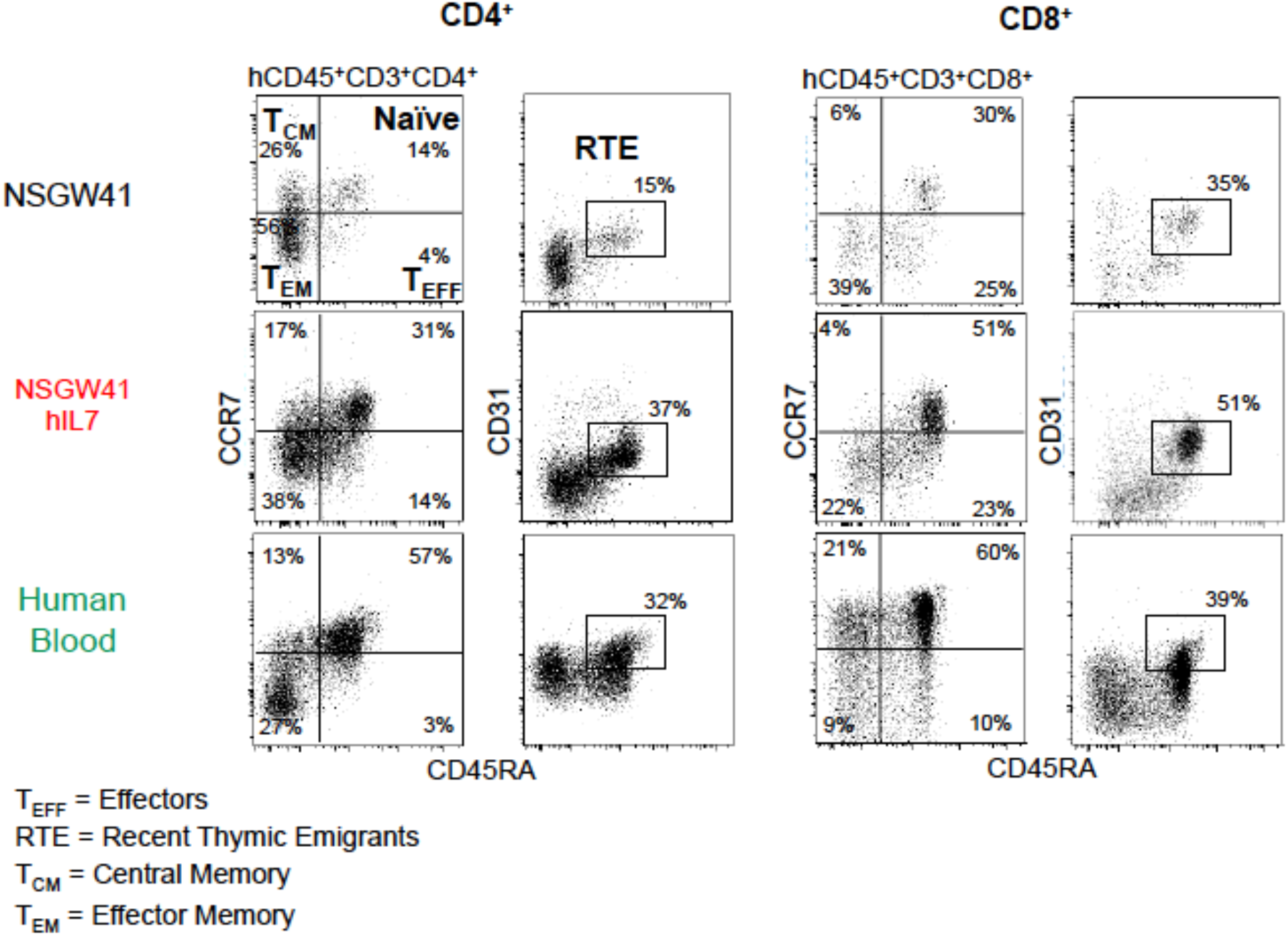
Characterization of human T cell subpopulations in the blood. Gating strategy and identification of hCD4^+^ and hCD8^+^ T cell subpopulations in blood of humanized mice: T effectors (T_EFF_, hCD45^+^ CD3^+^ CD4^+^ CCR7^−^ CD45RA^+^), central memory (T_CM_, hCD45^+^ CD3^+^ CD4^+^ CCR7^+^ CD45RA^−^), effector memory (T_EM_, hCD45^+^ CD3^+^ CD4^+^ CCR7^−^ CD45RA^−^), naïve T (Naïve, hCD45^+^ CD3^+^ CD4^+^ CCR7^+^ CD45RA^+^) and recent thymic emigrants (RTE, hCD45^+^ CD3^+^ CD4^+^ CD31^+^ CD45RA^+^). Data was acquired 26 weeks after humanization. The composition of subpopulations in human blood is shown for comparison (bottom).

**Supplementary Figure 4:**
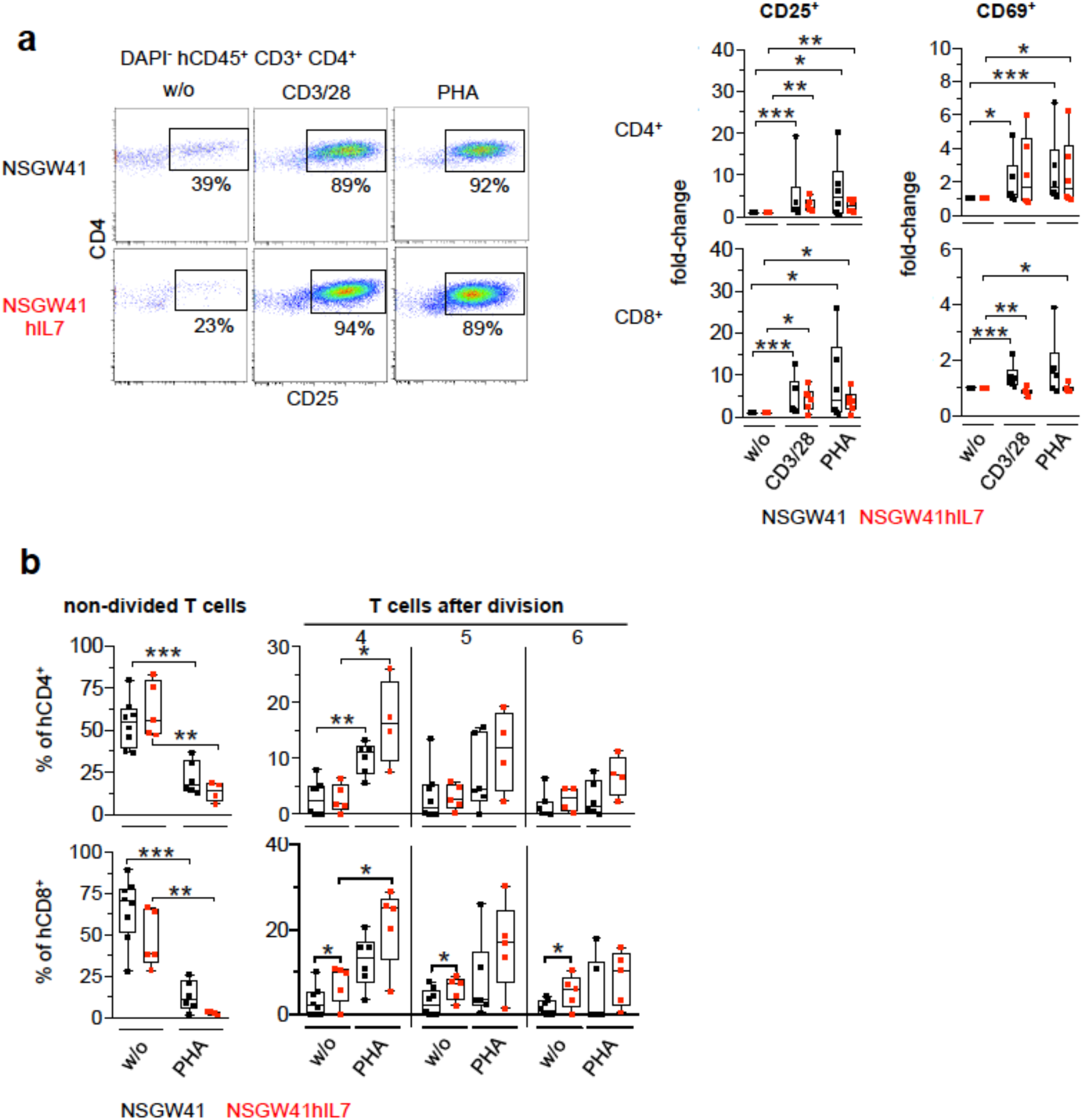
Activation of human T cells from NSGW41 or NSW41hIL7 mice. (**a**) Representative dot plots showing the expression of hCD25 and hCD69 on hCD3^+^ T cells isolated from NSGW41 (top) or NSGW41hIL7 (bottom) spleens after 6 days of CD3/28 or PHA stimulation or non-stimulated controls (left). Donor mice had received human HSPCs 20 to 26 weeks before. Fold-changes were calculated by dividing the percentages of stimulated hCD25^+^ or hCD69^+^ hCD4^+^ (top) or hCD8^+^ (bottom) splenic T cells by the percentages of non-stimulated cells (w/o) from NSGW41 or NSGW41hIL7 mice (right). (**b**) Percentages of non-divided T cells and T cells which have undergone 4, 5 or 6 divisions 6 days after stimulation with phytohemagglutinin (PHA) or without stimulation (w/o). Top: hCD4^+^ T cells, bottom: hCD8^+^ T cells. Donor mice had received human HSPCs 20 to 26 weeks before.

## Notes

### Competing Interest Statement

The authors have declared no competing interest.

